## Supporting material S1 for "Advancing science or advancing careers? Researchers’ opinions on success indicators"

### PRINTOUT OF THE SURVEY

Start of Block: Introduction

Welcome to our Survey for the project Re-SInC.

The survey aims to rethink research assessments and research careers. We want to know what **you** think.

You may find the full information for participation here.

To make this short, we thought we would give you three reasons for taking part in this survey!

- We need your input to know what really matters in research.
- It will only take 15-20 minutes of your time (we tested it with a few people, and it never took more than 16 minutes!).

You will help a very grateful PhD student graduate!

Note on ethics and privacy: This project has been approved by the Medical Ethics Committee of Hasselt University, protocol number CME2019/O35. Answers to the survey will be **fully confidential**, and **no identifiable information** (e.g., IP adresses, emails, etc.) is collected in the survey. The dataset, which contains **no identifiable information**, will be made public when findings are published.

Closing date: October 31st 2019
**If possible, use a computer to complete this questionnaire**.
If you only have a phone at hand, you should use the landscape mode.

End of Block: Introduction

Start of Block: Demographics

Before we start, we would like to get to know you!

You are a...

- o PhD Student
- o PostDoc / Non-tenure-track position
- o Tenure-track researcher / Professor
- o Tenured researcher / Full professor
- o I was a researcher in the past, but moved to another career
- o Other (specify) ________________________________________________

*Display This Question:*
*If Before we start, we would like to get to know you! You are a... = PhD Student*
*Or Before we start, we would like to get to know you! You are a... = PostDoc / Non-tenure-track position*
*Or Before we start, we would like to get to know you! You are a... = Tenure-track researcher / Professor*
*Or Before we start, we would like to get to know you! You are a... = Tenured researcher / Full professor*
*Or Before we start, we would like to get to know you! You are a... = Other (specify)*

You have been in this position for ...

▼ less than one year ... 9 years or more

*Display This Question:*
*If Before we start, we would like to get to know you! You are a... = I was a researcher in the past, but moved to another career*

How long ago did you stop being a researcher?

▼ less than one year ... 9 years or more

*Display This Question:*
*If Before we start, we would like to get to know you! You are a... = I was a researcher in the past, but moved to another career*

As someone who left academia your opinion is also very important to us! **Please answer the rest of this survey thinking back at your time as a researcher.**

Example: Read '***Are you*** *affiliated with a Flemish University or scientific institute?*' as '***Were you*** *affiliated with a Flemish University or scientific institute?*'

Are you affiliated with a Flemish University or scientific institute?

- Yes
- No

*Display This Question:*
*If Are you affiliated with a Flemish University or scientific institute? = Yes*

Your main affiliation is at...

- Hasselt University
- VU Brussels
- University of Antwerp
- Ghent University
- KU Leuven
- IMEC
- Institute of Tropical Medicine Antwerp
- Other ________________________________________________

*Display This Question:*
*If Are you affiliated with a Flemish University or scientific institute? = No*

Your main affiliation is at:

- Institution ________________________________________________
- Country ________________________________________________

You are in the Faculty of...

- Medicine / Medicine and Health Sciences / Medicine and Life Sciences / Medicine and Pharmacy *(or equivalent)*
- Other ________________________________________________

And you have published ...

▼ ...fewer than 10 peer-reviewed papers ... ...over 210 peer-reviewed papers

Your gender is...

- Male
- Female
- Other
- Prefer not to answer

Are you currently working in your country of origin?

- No
- Yes

*Display This Question:*
*If Are you currently working in your country of origin? = No*

What is your country of origin?

▼ Prefer not to say / Afghanistan ... Zimbabwe

*Display This Question:*
*If Are you currently working in your country of origin? = No*

Have you worked as a researcher/research student in your country of origin?

- Yes
- No

Have you ever been involved in **evaluating researchers for promotion, tenure, or career advancement**?

- Yes
- No
- Not sure (explain) ________________________________________________

**End of Block: Demographics**

**Start of Block: Time management**

Allright! Now we would like to know how you spend your time as a researcher.

Do you work full time as a researcher/PhD student?

- Yes
- No

*Display This Question:*
*If Allright! Now we would like to know how you spend your time as a researcher. Do you work full tim... = No*

Where would you situate yourself?

▼ less than 25% research employment ... over 75% but less than 100% research employment

On average, **HOW MANY HOURS** per week **do you work?** (overtime included)

I really work...

0 20 40 60 80

And during your work time, what **PERCENTAGE (%) of your time** do you spend on the following three pillars **in reality**, and **how would you like it to be**?

**In reality...**

**PERCENTAGE (%)** of your time

0 20 40 60 80 100

**TEACHING**
**RESEARCH**
**OTHER**

**In my ideal world...**

**PERCENTAGE (%)** of your time

0 20 40 60 80 100

**TEACHING**
**RESEARCH**
**OTHER**

And now a little more into the details:

What PERCENTAGE (%) of your time would you say you spend on average on each of the following activities **in reality**, and **how would you like it to be**?

**In reality...**

**PERCENTAGE (%)** of your time

0 20 40 60 80 100

**DIRECT STUDENT SUPERVISION**

**HANDS-ON RESEARCH WORK (e.g., lab work, data analysis)**

**STAYING UP TO DATE (e.g., reading, listening, building skills, etc.)**

**WRITING PAPERS**

**REVIEWING**

**GRANT WRITING**

**ANYTHING ELSE (e.g., administration, meetings, etc.)**

**In your ideal world...**

**PERCENTAGE (%)** of your time

0 20 40 60 80 100

**DIRECT STUDENT SUPERVISION**

**HANDS-ON RESEARCH WORK (e.g., lab work, data analysis)**

**STAYING UP TO DATE (e.g., reading, listening, building skills, etc.)**

**WRITING PAPERS**

**REVIEWING**

**GRANT WRITING**

**ANYTHING ELSE (e.g., administration, meetings, etc.)**

End of Block: Time management

---

Start of Block: Statements

Great! Now that we know each other, let's get to business!

In the following questions, we wish to know the impact of typical research activities
(A) on **advancing your career**
(B) on **advancing science**, and
(C) on **your personal satisfaction**

There will be **18 research activities** to rate. We numbered then from 18 to 1 so you know how many you have left!

**18. Publishing papers is...**

| | essential | important | irrelevant | unfavorable | detrimental |
|---|---|---|---|---|---|
| **...in advancing my career** | ○ | ○ | ○ | ○ | ○ |
| **...in advancing science** | ○ | ○ | ○ | ○ | ○ |
| **...to my personal satisfaction** | ○ | ○ | ○ | ○ | ○ |

Feel free to leave a comment (optional) ________________________________________________

**17. Publishing in high impact journals is...**

| | essential | important | irrelevant | unfavorable | detrimental |
|---|---|---|---|---|---|
| **...in advancing my career** | ○ | ○ | ○ | ○ | ○ |
| **...in advancing science** | ○ | ○ | ○ | ○ | ○ |
| **...to my personal satisfaction** | ○ | ○ | ○ | ○ | ○ |

Feel free to leave a comment (optional) ________________________________________________

**16. Publishing commentaries or editorials is…**

| | essential | important | irrelevant | unfavorable | detrimental |
|---|---|---|---|---|---|
| **…in advancing my career** | ○ | ○ | ○ | ○ | ○ |
| **…in advancing science** | ○ | ○ | ○ | ○ | ○ |
| **…to my personal satisfaction** | ○ | ○ | ○ | ○ | ○ |

Feel free to leave a comment (optional) ____________________________________________

**15. Publishing more papers than others is…**

| | essential | important | irrelevant | unfavorable | detrimental |
|---|---|---|---|---|---|
| **…in advancing my career** | ○ | ○ | ○ | ○ | ○ |
| **…in advancing science** | ○ | ○ | ○ | ○ | ○ |
| **…to my personal satisfaction** | ○ | ○ | ○ | ○ | ○ |

Feel free to leave a comment (optional) ____________________________________________

**14. Publishing open access is…**

| | essential | important | irrelevant | unfavorable | detrimental |
|---|---|---|---|---|---|
| **…in advancing my career** | ○ | ○ | ○ | ○ | ○ |
| **…in advancing science** | ○ | ○ | ○ | ○ | ○ |
| **…to my personal satisfaction** | ○ | ○ | ○ | ○ | ○ |

Feel free to leave a comment (optional) ____________________________________________

**13. Peer reviewing is…**

| | essential | important | irrelevant | unfavorable | detrimental |
|---|---|---|---|---|---|
| **…in advancing my career** | ○ | ○ | ○ | ○ | ○ |
| **…in advancing science** | ○ | ○ | ○ | ○ | ○ |
| **…to my personal satisfaction** | ○ | ○ | ○ | ○ | ○ |

Feel free to leave a comment (optional) ____________________________________________

**12. Replicating past research is…**

| | essential | important | irrelevant | unfavorable | detrimental |
|---|---|---|---|---|---|
| **…in advancing my career** | ○ | ○ | ○ | ○ | ○ |
| **…in advancing science** | ○ | ○ | ○ | ○ | ○ |
| **…to my personal satisfaction** | ○ | ○ | ○ | ○ | ○ |

Feel free to leave a comment (optional) ________________________________________________

**11. Publishing findings that did not work (i.e., negative findings) is…**

| | essential | important | irrelevant | unfavorable | detrimental |
|---|---|---|---|---|---|
| **…in advancing my career** | ○ | ○ | ○ | ○ | ○ |
| **…in advancing science** | ○ | ○ | ○ | ○ | ○ |
| **…to my personal satisfaction** | ○ | ○ | ○ | ○ | ○ |

Feel free to leave a comment (optional) ________________________________________________

**10. Sharing your full data and detailed methods is…**

| | essential | important | irrelevant | unfavorable | detrimental |
|---|---|---|---|---|---|
| **…in advancing my career** | ○ | ○ | ○ | ○ | ○ |
| **…in advancing science** | ○ | ○ | ○ | ○ | ○ |
| **…to my personal satisfaction** | ○ | ○ | ○ | ○ | ○ |

Feel free to leave a comment (optional) ________________________________________________

**9. Reviewing raw data from students and collaborators is…**

| | essential | important | irrelevant | unfavorable | detrimental |
|---|---|---|---|---|---|
| **…in advancing my career** | ○ | ○ | ○ | ○ | ○ |
| **…in advancing science** | ○ | ○ | ○ | ○ | ○ |
| **…to my personal satisfaction** | ○ | ○ | ○ | ○ | ○ |

Feel free to leave a comment (optional) ________________________________________________

**8. Conducting innovative research with a high risk of failure is...**

| | essential | important | irrelevant | unfavorable | detrimental |
|---|---|---|---|---|---|
| **...in advancing my career** | ○ | ○ | ○ | ○ | ○ |
| **...in advancing science** | ○ | ○ | ○ | ○ | ○ |
| **...to my personal satisfaction** | ○ | ○ | ○ | ○ | ○ |

Feel free to leave a comment (optional) ______________________________________________

**7. Connecting with renowned researchers is...**

| | essential | important | irrelevant | unfavorable | detrimental |
|---|---|---|---|---|---|
| **...in advancing my career** | ○ | ○ | ○ | ○ | ○ |
| **...in advancing science** | ○ | ○ | ○ | ○ | ○ |
| **...to my personal satisfaction** | ○ | ○ | ○ | ○ | ○ |

Feel free to leave a comment (optional) ______________________________________________

**6. Collaborating across borders, disciplines, and sectors is...**

| | essential | important | irrelevant | unfavorable | detrimental |
|---|---|---|---|---|---|
| **...in advancing my career** | ○ | ○ | ○ | ○ | ○ |
| **...in advancing science** | ○ | ○ | ○ | ○ | ○ |
| **...to my personal satisfaction** | ○ | ○ | ○ | ○ | ○ |

Feel free to leave a comment (optional) ______________________________________________

**5. Getting cited in scientific literature is...**

| | essential | important | irrelevant | unfavorable | detrimental |
|---|---|---|---|---|---|
| **...in advancing my career** | ○ | ○ | ○ | ○ | ○ |
| **...in advancing science** | ○ | ○ | ○ | ○ | ○ |
| **...to my personal satisfaction** | ○ | ○ | ○ | ○ | ○ |

Feel free to leave a comment (optional) ______________________________________________

**4. Having your papers read and downloaded is...**

| | essential | important | irrelevant | unfavorable | detrimental |
|---|---|---|---|---|---|
| **...in advancing my career** | ○ | ○ | ○ | ○ | ○ |
| **...in advancing science** | ○ | ○ | ○ | ○ | ○ |
| **...to my personal satisfaction** | ○ | ○ | ○ | ○ | ○ |

Feel free to leave a comment (optional) ________________________________________________

**3. Having public outreach (e.g., social media, news, etc.) is...**

| | essential | important | irrelevant | unfavorable | detrimental |
|---|---|---|---|---|---|
| **...in advancing my career** | ○ | ○ | ○ | ○ | ○ |
| **...in advancing science** | ○ | ○ | ○ | ○ | ○ |
| **...to my personal satisfaction** | ○ | ○ | ○ | ○ | ○ |

Feel free to leave a comment (optional) ________________________________________________

**2. Having your results used or implemented in practice is...**

| | essential | important | irrelevant | unfavorable | detrimental |
|---|---|---|---|---|---|
| **...in advancing my career** | ○ | ○ | ○ | ○ | ○ |
| **...in advancing science** | ○ | ○ | ○ | ○ | ○ |
| **...to my personal satisfaction** | ○ | ○ | ○ | ○ | ○ |

Feel free to leave a comment (optional) ________________________________________________

**1. Having luck is...**

| | essential | important | irrelevant | unfavorable | detrimental |
|---|---|---|---|---|---|
| **...in advancing my career** | ○ | ○ | ○ | ○ | ○ |
| **...in advancing science** | ○ | ○ | ○ | ○ | ○ |
| **...to my personal satisfaction** | ○ | ○ | ○ | ○ | ○ |

Feel free to leave a comment (optional) ________________________________________________

End of Block: Statements

Start of Block: End

Do you have any other comments or thoughts you would like to share with us? (optional)

________________________________________________________________
________________________________________________________________
__

End of Block: End
