## Supporting material S2 for "Advancing science or advancing careers? Researchers’ opinions on success indicators"

### CODE USED FOR THE ANALYSES

#### Time analyses in RStudio Version 1.1.453

```
# working directory
setwd("/Users/naubertbonn/RDirectorySurvey")
getwd()

#install.packages("dplyr")
library("dplyr")

FullDataset <- read.csv("S3.csv")

#Before anything, I need to make a dataset suited for comparing time distributions in 'Reality' and in an 'Ideal world'. This is a multi-step process that can be effectuated directly on the data file provided with the manuscript (Supplementary Data S3).

#Make a new dataset with the columns of interest (i.e., (1) ResponseID, (15) whether work full time as a researcher or not, and (18-37) all the answers on time distribution in reality and in an ideal world.)
TimeDetailsall <- FullDataset[c(1,15, 18:37)]

#And clean up the extra line that contain the full question text
TimeDetailsall <- TimeDetailsall[!(TimeDetailsall$ResponseId == "Response ID"),]

#Since this table was imported from the .csv, the data is not automatically considered in the right class, and we will need to convert some of the variables to numeric and character classes to allow our analyses.
#This first portion allows us to see which classes are applied to each variables
sapply (TimeDetailsall, class)

#At the moment, it is all classified under 'factor'. We need to have 'ResponseId', 'group', and 'FullTime' as character and all the rest as numeric.
TimeDetailsall$ResponseId <- as.character(TimeDetailsall$ResponseId)
TimeDetailsall$T1 <- as.character(TimeDetailsall$T1)
TimeDetailsall$T3_1 <- as.numeric(as.character(TimeDetailsall$T3_1))
TimeDetailsall$T3_2 <- as.numeric(as.character(TimeDetailsall$T3_2))
TimeDetailsall$T3_3 <- as.numeric(as.character(TimeDetailsall$T3_3))
TimeDetailsall$T4_1 <- as.numeric(as.character(TimeDetailsall$T4_1))
TimeDetailsall$T4_2 <- as.numeric(as.character(TimeDetailsall$T4_2))
TimeDetailsall$T4_3 <- as.numeric(as.character(TimeDetailsall$T4_3))
TimeDetailsall$T5_1 <- as.numeric(as.character(TimeDetailsall$T5_1))
TimeDetailsall$T5_2 <- as.numeric(as.character(TimeDetailsall$T5_2))
```

```
TimeDetailsall$T5_3 <- as.numeric(as.character(TimeDetailsall$T5_3))
TimeDetailsall$T5_4 <- as.numeric(as.character(TimeDetailsall$T5_4))
TimeDetailsall$T5_5 <- as.numeric(as.character(TimeDetailsall$T5_5))
TimeDetailsall$T5_6 <- as.numeric(as.character(TimeDetailsall$T5_6))
TimeDetailsall$T5_7 <- as.numeric(as.character(TimeDetailsall$T5_7))
TimeDetailsall$T6_1 <- as.numeric(as.character(TimeDetailsall$T6_1))
TimeDetailsall$T6_2 <- as.numeric(as.character(TimeDetailsall$T6_2))
TimeDetailsall$T6_3 <- as.numeric(as.character(TimeDetailsall$T6_3))
TimeDetailsall$T6_4 <- as.numeric(as.character(TimeDetailsall$T6_4))
TimeDetailsall$T6_5 <- as.numeric(as.character(TimeDetailsall$T6_5))
TimeDetailsall$T6_6 <- as.numeric(as.character(TimeDetailsall$T6_6))
TimeDetailsall$T6_7 <- as.numeric(as.character(TimeDetailsall$T6_7))


#Separate this dataset in two datasets (one for Reality and one ideal world) so we can make the values
 one on top of the other
TimeDetailsallReality <- TimeDetailsall[c(1, 2, 3:5, 9:15)]
TimeDetailsallIdeal <- TimeDetailsall[c(1,2, 6:8, 16:22)]
TimeDetailsallReality$group <- "Reality"
TimeDetailsallIdeal$group <- "Ideal"

#Rename the columns in each so that they correspond to each time category
colnames(TimeDetailsallIdeal)
names(TimeDetailsallReality)[names(TimeDetailsallReality) == "T1"] <- "FullTime"
names(TimeDetailsallIdeal)[names(TimeDetailsallIdeal) == "T1"] <- "FullTime"
names(TimeDetailsallReality)[names(TimeDetailsallReality) == "T3_1"] <- "Teaching"
names(TimeDetailsallIdeal)[names(TimeDetailsallIdeal) == "T4_1"] <- "Teaching"
names(TimeDetailsallReality)[names(TimeDetailsallReality) == "T3_2"] <- "Research"
names(TimeDetailsallIdeal)[names(TimeDetailsallIdeal) == "T4_2"] <- "Research"
names(TimeDetailsallReality)[names(TimeDetailsallReality) == "T3_3"] <- "Other"
names(TimeDetailsallIdeal)[names(TimeDetailsallIdeal) == "T4_3"] <- "Other"
names(TimeDetailsallReality)[names(TimeDetailsallReality) == "T5_1"] <- "Supervision"
names(TimeDetailsallIdeal)[names(TimeDetailsallIdeal) == "T6_1"] <- "Supervision"
names(TimeDetailsallReality)[names(TimeDetailsallReality) == "T5_2"] <- "ResearchWork"
names(TimeDetailsallIdeal)[names(TimeDetailsallIdeal) == "T6_2"] <- "ResearchWork"
names(TimeDetailsallReality)[names(TimeDetailsallReality) == "T5_3"] <- "UpToDate"
names(TimeDetailsallIdeal)[names(TimeDetailsallIdeal) == "T6_3"] <- "UpToDate"
names(TimeDetailsallReality)[names(TimeDetailsallReality) == "T5_4"] <- "Writing"
names(TimeDetailsallIdeal)[names(TimeDetailsallIdeal) == "T6_4"] <- "Writing"
names(TimeDetailsallReality)[names(TimeDetailsallReality) == "T5_5"] <- "Reviewing"
names(TimeDetailsallIdeal)[names(TimeDetailsallIdeal) == "T6_5"] <- "Reviewing"
names(TimeDetailsallReality)[names(TimeDetailsallReality) == "T5_6"] <- "GrantWriting"
names(TimeDetailsallIdeal)[names(TimeDetailsallIdeal) == "T6_6"] <- "GrantWriting"
names(TimeDetailsallReality)[names(TimeDetailsallReality) == "T5_7"] <- "AnythingElse"
names(TimeDetailsallIdeal)[names(TimeDetailsallIdeal) == "T6_7"] <- "AnythingElse"
colnames(TimeDetailsallIdeal)
```

```
#Merge the two sets so that we have all 'Reality' and 'Ideal' Times in the same dataset
TimeDetailsall <- rbind(TimeDetailsallReality, TimeDetailsallIdeal)

#Keep only full-time researchers  in a separate set to see if their answers differ from non-full-time
  researcher respondents
TimeDetails <- TimeDetailsall[ which(TimeDetailsall$FullTime=='Yes'), ]

#Compute the log of all, just to be able to compare the answers of those working full time to those
  working part time
TimeDetailsall$LogTeachingall <- log10(TimeDetailsall$Teaching+0.5)
TimeDetailsall$LogResearchall <- log10(TimeDetailsall$Research+0.5)
TimeDetailsall$LogOtherall <- log10(TimeDetailsall$Other+0.5)
TimeDetailsall$LogSupervisionall <- log10(TimeDetailsall$Supervision+0.5)
TimeDetailsall$LogResearchWorkall <- log10(TimeDetailsall$ResearchWork+0.5)
TimeDetailsall$LogUpToDateall <- log10(TimeDetailsall$UpToDate+0.5)
TimeDetailsall$LogWritingall <- log10(TimeDetailsall$Writing+0.5)
TimeDetailsall$LogReviewingall <- log10(TimeDetailsall$Reviewing+0.5)
TimeDetailsall$LogGrantWritingall <- log10(TimeDetailsall$GrantWriting+0.5)
TimeDetailsall$LogAnythingElseall <- log10(TimeDetailsall$AnythingElse+0.5)

#For each time category, test whether the groups (full-time/non-full-time) are different
TTTeachingall <- t.test(LogTeachingall ~ FullTime, data = TimeDetailsall, paired = FALSE)
TTResearchall <- t.test(LogResearchall ~ FullTime, data = TimeDetailsall, paired = FALSE)
TTOtherall <- t.test(LogOtherall ~ FullTime, data = TimeDetailsall, paired = FALSE)
TTSupervisionall <- t.test(LogSupervisionall ~ FullTime, data = TimeDetailsall, paired = FALSE)
TTResearchWorkall <- t.test(LogResearchWorkall ~ FullTime, data = TimeDetailsall, paired = FALSE)
TTUpToDateall <- t.test(LogUpToDateall ~ FullTime, data = TimeDetailsall, paired = FALSE)
TTWritingall <- t.test(LogWritingall ~ FullTime, data = TimeDetailsall, paired = FALSE)
TTReviewingall <- t.test(LogReviewingall ~ FullTime, data = TimeDetailsall, paired = FALSE)
TTGrantWritingall <- t.test(LogGrantWritingall ~ FullTime, data = TimeDetailsall, paired = FALSE)
TTAnythingElseall <- t.test(LogAnythingElseall ~ FullTime, data = TimeDetailsall, paired = FALSE)

TTTeachingall
TTResearchall
TTOtherall
TTSupervisionall
TTResearchWorkall
TTUpToDateall
TTWritingall
TTReviewingall
TTGrantWritingall
TTAnythingElseall

# RESULTS ---> 'Research', 'Other' / 'Hands-on Research work'and 'AnythingElse' differed between full
  time and part time respondents. As a result, I chose to keep only full time respondents for the next
  analyses...
```

```
#Test normality
#After discussing with the Editor of this paper, we decided that the normality assumption was not
 relevant
#with our data type (i.e., percentages) and therefore analysed the data as is. Former code and analyses
 are
#available in older versions of our preprint at https://doi.org/10.1101/2020.06.22.165654

# Paired t-test to compare respondents' time distributions in 'reality' to how they which they could
 distribute their time in an 'ideal' world
TTTeaching <- t.test(Teaching ~ group, data = TimeDetails, paired = TRUE)
TTResearch <- t.test(Research ~ group, data = TimeDetails, paired = TRUE)
TTOther <- t.test(Other ~ group, data = TimeDetails, paired = TRUE)
TTSupervision <- t.test(Supervision ~ group, data = TimeDetails, paired = TRUE)
TTResearchWork <- t.test(ResearchWork ~ group, data = TimeDetails, paired = TRUE)
TTUpToDate <- t.test(UpToDate ~ group, data = TimeDetails, paired = TRUE)
TTWriting <- t.test(Writing ~ group, data = TimeDetails, paired = TRUE)
TTReviewing <- t.test(Reviewing ~ group, data = TimeDetails, paired = TRUE)
TTGrantWriting <- t.test(GrantWriting ~ group, data = TimeDetails, paired = TRUE)
TTAnythingElse <- t.test(AnythingElse ~ group, data = TimeDetails, paired = TRUE)


#And display the result first of the T Test, and then the measures of central tendencies related to the
 relevant dimension

#Display the T Test Result
TTTeaching
#And display results of the mean, median, and SD
library("dplyr")
group_by(TimeDetails, group) %>%
  summarise(
    count = n(),
    mean = mean(Teaching, na.rm = TRUE),
    median = median(Teaching, na.rm = TRUE),
    sd = sd(Teaching, na.rm = TRUE)
  )

#Display the T Test Result
TTResearch
#And display results of the mean, median, and SD
library("dplyr")
group_by(TimeDetails, group) %>%
  summarise(
    count = n(),
    mean = mean(Research, na.rm = TRUE),
    median = median(Research, na.rm = TRUE),
    sd = sd(Research, na.rm = TRUE)
  )
```

```
#Display the T Test Result
TTOther
#And display results of the mean, median, and SD
library("dplyr")
group_by(TimeDetails, group) %>%
  summarise(
    count = n(),
    mean = mean(Other, na.rm = TRUE),
    median = median(Other, na.rm = TRUE),
    sd = sd(Other, na.rm = TRUE)
  )

#Display the T Test Result
TTSupervision
#And display results of the mean, median, and SD
library("dplyr")
group_by(TimeDetails, group) %>%
  summarise(
    count = n(),
    mean = mean(Supervision, na.rm = TRUE),
    median = median(Supervision, na.rm = TRUE),
    sd = sd(Supervision, na.rm = TRUE)
  )

#Display the T Test Result
TTResearchWork
#And display results of the mean, median, and SD
library("dplyr")
group_by(TimeDetails, group) %>%
  summarise(
    count = n(),
    mean = mean(ResearchWork, na.rm = TRUE),
    median = median(ResearchWork, na.rm = TRUE),
    sd = sd(ResearchWork, na.rm = TRUE)
  )

#Display the T Test Result
TTUpToDate
#And display results of the mean, median, and SD
library("dplyr")
group_by(TimeDetails, group) %>%
  summarise(
    count = n(),
    mean = mean(UpToDate, na.rm = TRUE),
    median = median(UpToDate, na.rm = TRUE),
    sd = sd(UpToDate, na.rm = TRUE)
```

```
  )

#Display the T Test Result
TTWriting
#And display results of the mean, median, and SD
library("dplyr")
group_by(TimeDetails, group) %>%
  summarise(
    count = n(),
    mean = mean(Writing, na.rm = TRUE),
    median = median(Writing, na.rm = TRUE),
    sd = sd(Writing, na.rm = TRUE)
  )

#Display the T Test Result
TTReviewing
#And display results of the mean, median, and SD
library("dplyr")
group_by(TimeDetails, group) %>%
  summarise(
    count = n(),
    mean = mean(Reviewing, na.rm = TRUE),
    median = median(Reviewing, na.rm = TRUE),
    sd = sd(Reviewing, na.rm = TRUE)
  )

#Display the T Test Result
TTGrantWriting
#And display results of the mean, median, and SD
library("dplyr")
group_by(TimeDetails, group) %>%
  summarise(
    count = n(),
    mean = mean(GrantWriting, na.rm = TRUE),
    median = median(GrantWriting, na.rm = TRUE),
    sd = sd(GrantWriting, na.rm = TRUE)
  )

#Display the T Test Result
TTAnythingElse
#And display results of the mean, median, and SD
library("dplyr")
group_by(TimeDetails, group) %>%
  summarise(
    count = n(),
    mean = mean(AnythingElse, na.rm = TRUE),
    median = median(AnythingElse, na.rm = TRUE),
```

```
  sd = sd(AnythingElse, na.rm = TRUE)
 )
```

### Dimension analyses in IBM® SPSS® Statistics version 25

Note: The following code is the code corresponding to the SPSS analyses. The file used to perform these analyses can be automatically generates in R using the code copied at the end of this Supplementary file.

```
GLM S1Career S1Science S1Satisfaction
  /WSFACTOR=Dimension 3 Polynomial
  /MEASURE=importance
  /METHOD=SSTYPE(3)
  /EMMEANS=TABLES(Dimension) COMPARE ADJ(LSD)
  /PRINT=DESCRIPTIVE
  /CRITERIA=ALPHA(.05)
  /WSDESIGN=Dimension.

GLM S2Career S2Science S2Satisfaction
  /WSFACTOR=Dimension 3 Polynomial
  /MEASURE=importance
  /METHOD=SSTYPE(3)
  /EMMEANS=TABLES(Dimension) COMPARE ADJ(LSD)
  /PRINT=DESCRIPTIVE
  /CRITERIA=ALPHA(.05)
  /WSDESIGN=Dimension.

GLM S3Career S3Science S3Satisfaction
  /WSFACTOR=Dimension 3 Polynomial
  /MEASURE=importance
  /METHOD=SSTYPE(3)
  /EMMEANS=TABLES(Dimension) COMPARE ADJ(LSD)
  /PRINT=DESCRIPTIVE
  /CRITERIA=ALPHA(.05)
  /WSDESIGN=Dimension.

GLM S4Career S4Science S4Satisfaction
  /WSFACTOR=Dimension 3 Polynomial
  /MEASURE=importance
  /METHOD=SSTYPE(3)
  /EMMEANS=TABLES(Dimension) COMPARE ADJ(LSD)
  /PRINT=DESCRIPTIVE
  /CRITERIA=ALPHA(.05)
  /WSDESIGN=Dimension.

GLM S5Career S5Science S5Satisfaction
  /WSFACTOR=Dimension 3 Polynomial
```

```
 /MEASURE=importance
 /METHOD=SSTYPE(3)
 /EMMEANS=TABLES(Dimension) COMPARE ADJ(LSD)
 /PRINT=DESCRIPTIVE
 /CRITERIA=ALPHA(.05)
 /WSDESIGN=Dimension.

GLM S6Career S6Science S6Satisfaction
 /WSFACTOR=Dimension 3 Polynomial
 /MEASURE=importance
 /METHOD=SSTYPE(3)
 /EMMEANS=TABLES(Dimension) COMPARE ADJ(LSD)
 /PRINT=DESCRIPTIVE
 /CRITERIA=ALPHA(.05)
 /WSDESIGN=Dimension.

GLM S7Career S7Science S7Satisfaction
 /WSFACTOR=Dimension 3 Polynomial
 /MEASURE=importance
 /METHOD=SSTYPE(3)
 /EMMEANS=TABLES(Dimension) COMPARE ADJ(LSD)
 /PRINT=DESCRIPTIVE
 /CRITERIA=ALPHA(.05)
 /WSDESIGN=Dimension.

GLM S8Career S8Science S8Satisfaction
 /WSFACTOR=Dimension 3 Polynomial
 /MEASURE=importance
 /METHOD=SSTYPE(3)
 /EMMEANS=TABLES(Dimension) COMPARE ADJ(LSD)
 /PRINT=DESCRIPTIVE
 /CRITERIA=ALPHA(.05)
 /WSDESIGN=Dimension.

GLM S9Career S9Science S9Satisfaction
 /WSFACTOR=Dimension 3 Polynomial
 /MEASURE=importance
 /METHOD=SSTYPE(3)
 /EMMEANS=TABLES(Dimension) COMPARE ADJ(LSD)
 /PRINT=DESCRIPTIVE
 /CRITERIA=ALPHA(.05)
 /WSDESIGN=Dimension.

GLM S10Career S10Science S10Satisfaction
 /WSFACTOR=Dimension 3 Polynomial
 /MEASURE=importance
 /METHOD=SSTYPE(3)
```

```
 /EMMEANS=TABLES(Dimension) COMPARE ADJ(LSD)
 /PRINT=DESCRIPTIVE
 /CRITERIA=ALPHA(.05)
 /WSDESIGN=Dimension.

GLM S11Career S11Science S11Satisfaction
 /WSFACTOR=Dimension 3 Polynomial
 /MEASURE=importance
 /METHOD=SSTYPE(3)
 /EMMEANS=TABLES(Dimension) COMPARE ADJ(LSD)
 /PRINT=DESCRIPTIVE
 /CRITERIA=ALPHA(.05)
 /WSDESIGN=Dimension.

GLM S12Career S12Science S12Satisfaction
 /WSFACTOR=Dimension 3 Polynomial
 /MEASURE=importance
 /METHOD=SSTYPE(3)
 /EMMEANS=TABLES(Dimension) COMPARE ADJ(LSD)
 /PRINT=DESCRIPTIVE
 /CRITERIA=ALPHA(.05)
 /WSDESIGN=Dimension.

GLM S13Career S13Science S13Satisfaction
 /WSFACTOR=Dimension 3 Polynomial
 /MEASURE=importance
 /METHOD=SSTYPE(3)
 /EMMEANS=TABLES(Dimension) COMPARE ADJ(LSD)
 /PRINT=DESCRIPTIVE
 /CRITERIA=ALPHA(.05)
 /WSDESIGN=Dimension.

GLM S14Career S14Science S14Satisfaction
 /WSFACTOR=Dimension 3 Polynomial
 /MEASURE=importance
 /METHOD=SSTYPE(3)
 /EMMEANS=TABLES(Dimension) COMPARE ADJ(LSD)
 /PRINT=DESCRIPTIVE
 /CRITERIA=ALPHA(.05)
 /WSDESIGN=Dimension.

GLM S15Career S15Science S15Satisfaction
 /WSFACTOR=Dimension 3 Polynomial
 /MEASURE=importance
 /METHOD=SSTYPE(3)
 /EMMEANS=TABLES(Dimension) COMPARE ADJ(LSD)
 /PRINT=DESCRIPTIVE
```

```
 /CRITERIA=ALPHA(.05)
 /WSDESIGN=Dimension.

GLM S16Career S16Science S16Satisfaction
 /WSFACTOR=Dimension 3 Polynomial
 /MEASURE=importance
 /METHOD=SSTYPE(3)
 /EMMEANS=TABLES(Dimension) COMPARE ADJ(LSD)
 /PRINT=DESCRIPTIVE
 /CRITERIA=ALPHA(.05)
 /WSDESIGN=Dimension.

GLM S17Career S17Science S17Satisfaction
 /WSFACTOR=Dimension 3 Polynomial
 /MEASURE=importance
 /METHOD=SSTYPE(3)
 /EMMEANS=TABLES(Dimension) COMPARE ADJ(LSD)
 /PRINT=DESCRIPTIVE
 /CRITERIA=ALPHA(.05)
 /WSDESIGN=Dimension.

GLM S18Career S18Science S18Satisfaction
 /WSFACTOR=Dimension 3 Polynomial
 /MEASURE=importance
 /METHOD=SSTYPE(3)
 /EMMEANS=TABLES(Dimension) COMPARE ADJ(LSD)
 /PRINT=DESCRIPTIVE
 /CRITERIA=ALPHA(.05)
 /WSDESIGN=Dimension.


GET DATA
 /TYPE=XLSX
 /FILE='/Volumes/…/Survey/Data/SPSS/QuestionsStatements.xlsx'
 /SHEET=name 'Responses Statements'
 /CELLRANGE=FULL
 /READNAMES=ON
 /DATATYPEMIN PERCENTAGE=95.0
 /HIDDEN IGNORE=YES.
EXECUTE.
DATASET NAME DataSet5 WINDOW=FRONT.
DATASET CLOSE DataSet4.
DESCRIPTIVES VARIABLES=S1Career S1Science S1Satisfaction S2Career S2Science S2Satisfaction
S3Career
   S3Science S3Satisfaction S4Career S4Science S4Satisfaction S5Career S5Science S5Satisfaction
   S6Career S6Science S6Satisfaction S7Career S7Science S7Satisfaction S8Career S8Science
   S8Satisfaction S9Career S9Science S9Satisfaction S10Career S10Science S10Satisfaction S11Career
```

```
   S11Science S11Satisfaction S12Career S12Science S12Satisfaction S13Career S13Science
   S13Satisfaction S14Career S14Science S14Satisfaction S15Career S15Science S15Satisfaction
S16Career
   S16Science S16Satisfaction S17Career S17Science S17Satisfaction S18Career S18Science
S18Satisfaction
  /STATISTICS=MEAN STDDEV MIN MAX.
```

**R code which can be used to generate the dataset for use in SPSS dimension analysis**

```
#This code is run in R simply to generate the dataset for the SPSS analyses

# working directory
setwd("/Users/naubertbonn/RDirectorySurvey")
getwd()

#install.packages("dplyr")
#install.packages("xlsx")
library("dplyr")

FullDataset <- read.csv("S3.csv")

#Make a new dataset with the columns (1) ResponseID, and all columns about the ratings of
  success indicators(15) whether work full time as a researcher or not, and (18-37) all the
  answers on time distribution in reality and in an ideal world.)
SuccessIndicators <- FullDataset[c(1, 38:109)]
SuccessIndicators <- subset(SuccessIndicators, select=-
  c(5,9,13,17,21,25,29,33,37,41,45,49,53,57,61,65,69,73)) #to drop the comments columns
colnames(SuccessIndicators)

#And clean up the extra line that contain the full question text
SuccessIndicators <- SuccessIndicators[!(SuccessIndicators$ResponseId == "Response ID"),]

#Since this table was imported from the .csv, the data is not automatically considered in the
  right class, and we will need to convert some of the variables to numeric and character
  classes to allow our analyses.
#This first portion allows us to see which classes are applied to each variables
sapply (SuccessIndicators, class)

#At the moment, it is all classified under 'factor'. We need to have 'ResponseId', 'group', and
  'FullTime' as character and all the rest as numeric.
SuccessIndicators$ResponseId <- as.character(SuccessIndicators$ResponseId)
SuccessIndicators[,2:55] <- lapply(SuccessIndicators[,2:55], as.character)

#Before transforming to numeric variables, we need to change the values from the string to
  the numbers
SuccessIndicators [SuccessIndicators == "essential"] <- "5"
SuccessIndicators [SuccessIndicators == "important"] <- "4"
```

```
SuccessIndicators [SuccessIndicators == "irrelevant"] <- "3"
SuccessIndicators [SuccessIndicators == "unfavorable"] <- "2"
SuccessIndicators [SuccessIndicators == "detrimental"] <- "1"

#Then we transform all these character variables to numeric variables
SuccessIndicators[,2:55] <- lapply(SuccessIndicators[,2:55], as.numeric)

#And for simplicity of interpretation, we can finally change the column names so we
  understand which dimention they represent
names(SuccessIndicators)[names(SuccessIndicators) == "S1_1"] <- "S1Career"
names(SuccessIndicators)[names(SuccessIndicators) == "S1_2"] <- "S1Science"
names(SuccessIndicators)[names(SuccessIndicators) == "S1_3"] <- "S1Satisfaction"
names(SuccessIndicators)[names(SuccessIndicators) == "S2_1"] <- "S2Career"
names(SuccessIndicators)[names(SuccessIndicators) == "S2_2"] <- "S2Science"
names(SuccessIndicators)[names(SuccessIndicators) == "S2_3"] <- "S2Satisfaction"
names(SuccessIndicators)[names(SuccessIndicators) == "S3_1"] <- "S3Career"
names(SuccessIndicators)[names(SuccessIndicators) == "S3_2"] <- "S3Science"
names(SuccessIndicators)[names(SuccessIndicators) == "S3_3"] <- "S3Satisfaction"
names(SuccessIndicators)[names(SuccessIndicators) == "S4_1"] <- "S4Career"
names(SuccessIndicators)[names(SuccessIndicators) == "S4_2"] <- "S4Science"
names(SuccessIndicators)[names(SuccessIndicators) == "S4_3"] <- "S4Satisfaction"
names(SuccessIndicators)[names(SuccessIndicators) == "S5_1"] <- "S5Career"
names(SuccessIndicators)[names(SuccessIndicators) == "S5_2"] <- "S5Science"
names(SuccessIndicators)[names(SuccessIndicators) == "S5_3"] <- "S5Satisfaction"
names(SuccessIndicators)[names(SuccessIndicators) == "S6_1"] <- "S6Career"
names(SuccessIndicators)[names(SuccessIndicators) == "S6_2"] <- "S6Science"
names(SuccessIndicators)[names(SuccessIndicators) == "S6_3"] <- "S6Satisfaction"
names(SuccessIndicators)[names(SuccessIndicators) == "S7_1"] <- "S7Career"
names(SuccessIndicators)[names(SuccessIndicators) == "S7_2"] <- "S7Science"
names(SuccessIndicators)[names(SuccessIndicators) == "S7_3"] <- "S7Satisfaction"
names(SuccessIndicators)[names(SuccessIndicators) == "S8_1"] <- "S8Career"
names(SuccessIndicators)[names(SuccessIndicators) == "S8_2"] <- "S8Science"
names(SuccessIndicators)[names(SuccessIndicators) == "S8_3"] <- "S8Satisfaction"
names(SuccessIndicators)[names(SuccessIndicators) == "S9_1"] <- "S9Career"
names(SuccessIndicators)[names(SuccessIndicators) == "S9_2"] <- "S9Science"
names(SuccessIndicators)[names(SuccessIndicators) == "S9_3"] <- "S9Satisfaction"
names(SuccessIndicators)[names(SuccessIndicators) == "S10_1"] <- "S10Career"
names(SuccessIndicators)[names(SuccessIndicators) == "S10_2"] <- "S10Science"
names(SuccessIndicators)[names(SuccessIndicators) == "S10_3"] <- "S10Satisfaction"
names(SuccessIndicators)[names(SuccessIndicators) == "S11_1"] <- "S11Career"
names(SuccessIndicators)[names(SuccessIndicators) == "S11_2"] <- "S11Science"
names(SuccessIndicators)[names(SuccessIndicators) == "S11_3"] <- "S11Satisfaction"
names(SuccessIndicators)[names(SuccessIndicators) == "S12_1"] <- "S12Career"
names(SuccessIndicators)[names(SuccessIndicators) == "S12_2"] <- "S12Science"
names(SuccessIndicators)[names(SuccessIndicators) == "S12_3"] <- "S12Satisfaction"
names(SuccessIndicators)[names(SuccessIndicators) == "S13_1"] <- "S13Career"
names(SuccessIndicators)[names(SuccessIndicators) == "S13_2"] <- "S13Science"
```

```
names(SuccessIndicators)[names(SuccessIndicators) == "S13_3"] <- "S13Satisfaction"
names(SuccessIndicators)[names(SuccessIndicators) == "S14_1"] <- "S14Career"
names(SuccessIndicators)[names(SuccessIndicators) == "S14_2"] <- "S14Science"
names(SuccessIndicators)[names(SuccessIndicators) == "S14_3"] <- "S14Satisfaction"
names(SuccessIndicators)[names(SuccessIndicators) == "S15_1"] <- "S15Career"
names(SuccessIndicators)[names(SuccessIndicators) == "S15_2"] <- "S15Science"
names(SuccessIndicators)[names(SuccessIndicators) == "S15_3"] <- "S15Satisfaction"
names(SuccessIndicators)[names(SuccessIndicators) == "S16_1"] <- "S16Career"
names(SuccessIndicators)[names(SuccessIndicators) == "S16_2"] <- "S16Science"
names(SuccessIndicators)[names(SuccessIndicators) == "S16_3"] <- "S16Satisfaction"
names(SuccessIndicators)[names(SuccessIndicators) == "S17_1"] <- "S17Career"
names(SuccessIndicators)[names(SuccessIndicators) == "S17_2"] <- "S17Science"
names(SuccessIndicators)[names(SuccessIndicators) == "S17_3"] <- "S17Satisfaction"
names(SuccessIndicators)[names(SuccessIndicators) == "S18_1"] <- "S18Career"
names(SuccessIndicators)[names(SuccessIndicators) == "S18_2"] <- "S18Science"
names(SuccessIndicators)[names(SuccessIndicators) == "S18_3"] <- "S18Satisfaction"

#And finally, export the results in an Excel file to be used in SPSS. This file will be saved in
 your R Directory, which will be displayed thereafter
getwd()
library("xlsx")
write.csv(SuccessIndicators, file="DimensionsDataForSPSS.xls", sheetName = "Sheet1",
        col.names = TRUE, row.names = FALSE, append = FALSE)
```
