## Supporting table S4 for "Advancing science or advancing careers? Researchers’ opinions on success indicators"

### SUPPLEMENTARY TABLE S4

#### ANALYSIS RESULTS FOR RATINGS OF THE DIMENSIONAL IMPORTANCE OF THE 18 SUCCESS INDICATORS

| | | | Bonferroni post hoc pairwise comparisons | | |
|---|---|---|---|---|---|
| Statement | Mean rating for each dimension | Result* | Career vs Science | Career vs Satisfaction | Science vs Satisfaction |
| Publishing papers is... | Career: 4.52<br>Science: 4.37<br>Satisfaction: 3.90 | (SA) F(2, 250) = 38.188<br>p < 0.001 | 95% CI(0.001, 0.301)<br>p = 0.048<br>md = 0.151 | **95% CI(0.468, 0.786)<br>p < 0.001<br>md = 0.627** | **95% CI(0.341, 0.611)<br>p < 0.001<br>md = 0.476** |
| Publishing in high impact journals is... | Career: 4.31<br>Science: 3.73<br>Satisfaction: 3.67 | (SA) F(2, 250) = 36.701<br>p < 0.001 | **95% CI(0.418, 0.741)<br>p < 0.001<br>md = 0.579** | **95% CI(0.477, 0.809)<br>p < 0.001<br>md = 0.643** | 95% CI(−0.100, 0.227)<br>p = 0.444<br>md = 0.063 |
| Publishing commentaries or editorials is... | Career: 3.48<br>Science: 3.70<br>Satisfaction: 3.34 | (SA) F(2, 250) = 14.538<br>p < 0.001 | **95% CI(−0.360, −0.085)<br>p = 0.002<br>md = −0.222** | 95% CI(0.000, 0.269)<br>p = 0.049<br>md = 0.135 | **95% CI(0.232, 0.482)<br>p < 0.001<br>md = 0.357** |
| Publishing more papers than others is... | Career: 3.83<br>Science: 2.89<br>Satisfaction: 3.01 | (GG) F(1.779, 222.388)<br>= 70.233<br>p < 0.001 | **95% CI(0.761, 1.127)<br>p < 0.001<br>md = 0.944** | **95% CI(0.636, 1.015)<br>p < 0.001<br>md = 0.825** | 95% CI(−0.258, 0.020)<br>p = 0.092<br>md = −0.119 |
| Publishing open access is... | Career: 3.48<br>Science: 4.35<br>Satisfaction: 3.67 | (SA) F(2, 250) = 62.624<br>p < 0.001 | **95% CI(−1.034, −0.696)<br>p < 0.001<br>md = −0.865** | **95% CI(−0.357, −0.024)<br>p = 0.025<br>md = −0.190** | **95% CI(0.529, 0.821)<br>p < 0.001<br>md = 0.675** |
| Peer reviewing is... | Career: 3.39<br>Science: 4.43<br>Satisfaction: 3.47 | (SA) F(2, 250) = 81.399<br>p < 0.001 | **95% CI(−1.221, −0.858)<br>p < 0.001<br>md = −1.040** | 95% CI(−0.260, 0.101)<br>p = 0.386<br>md = −0.079 | **95% CI(0.784, 1.136)<br>p < 0.001<br>md = 0.960** |
| Replicating past research is... | Career: 2.83<br>Science: 3.98<br>Satisfaction: 3.09 | (GG) F(1.847, 230.832)<br>= 81.530<br>p < 0.001 | **95% CI(−1.360, −0.942)<br>p < 0.001<br>md = −1.151** | **95% CI(−0.442, −0.066)<br>p = 0.008<br>md = −0.254** | **95% CI(0.735, 1.059)<br>p < 0.001<br>md = 0.897** |
| Publishing findings that did not work (i.e., negative findings) is... | Career: 2.84<br>Science: 4.48<br>Satisfaction: 3.61 | (GG) F(1.820, 227.487)<br>= 187.113<br>p < 0.001 | **95% CI(−1.823, −1.462)<br>p < 0.001<br>md = −1.643** | **95% CI(−0.951, −0.588)<br>p < 0.001<br>md = −0.770** | **95% CI(0.734, 1.012)<br>p < 0.001<br>md = 0.873** |
| Sharing your full data and detailed methods is... | Career: 3.29<br>Science: 4.40<br>Satisfaction: 3.67 | (GG) F(1.906, 238.310)<br>= 106.656<br>p < 0.001 | **95% CI(−1.278, −0.960)<br>p < 0.001<br>md = −1.119** | **95% CI(−0.554, −0.224)<br>p < 0.001<br>md = −0.389** | **95% CI(0.594, 0.867)<br>p < 0.001<br>md = 0.730** |
| Reviewing raw data from students and collaborators is... | Career: 3.37<br>Science: 4.20<br>Satisfaction: 3.64 | (SA) F(2, 250) = 50.707<br>p < 0.001 | **95% CI(−0.992, −0.658)<br>p < 0.001<br>md = −0.825** | **95% CI(−0.446, −0.094)<br>p = 0.003<br>md = −0.270** | **95% CI(0.403, 0.708)<br>p < 0.001<br>md = 0.556** |
| Conducting innovative research with a high risks of failure is... | Career: 3.29<br>Science: 4.47<br>Satisfaction: 3.91 | (GG) F(1.614, 201.766)<br>= 66.452<br>p < 0.001 | **95% CI(−1.406, −0.943)<br>p < 0.001<br>md = −1.175** | **95% CI(−0.836, −0.402)<br>p < 0.001<br>md = −0.619** | **95% CI(0.410, 0.701)<br>p < 0.001<br>md = 0.556** |
| Connecting with renowned researchers is... | Career: 4.35<br>Science: 3.91<br>Satisfaction: 3.98 | (SA) F(2, 250) = 24.566<br>p < 0.001 | **95% CI(0.295, 0.578)<br>p < 0.001<br>md = 0.437** | **95% CI(0.238, 0.508)<br>p < 0.001<br>md = 0.373** | 95% CI(−0.185, 0.058)<br>p = 0.304<br>md = −0.063 |
| Collaborating across borders, disciplines, and sectors is... | Career: 4.25<br>Science: 4.64<br>Satisfaction: 4.36 | (GG) F(1.558, 194.781)<br>= 14.565<br>p < 0.001 | **95% CI(−0.550, −0.228)<br>p < 0.001<br>md = −0.389** | 95% CI(−0.274, 0.068)<br>p = 0.235<br>md = −0.103 | **95% CI(0.184, 0.388)<br>p < 0.001<br>md = 0.286** |
| Getting cited in scientific literature is... | Career: 4.46<br>Science: 3.66<br>Satisfaction: 3.98 | (GG) F(1.890, 236.280)<br>= 55.630<br>p < 0.001 | **95% CI(0.633, 0.970)<br>p < 0.001<br>md = 0.802** | **95% CI(0.345, 0.623)<br>p < 0.001<br>md = 0.484** | **95% CI(−0.463, −0.172)<br>p < 0.001<br>md = −0.317** |

| | | | | | |
|---|---|---|---|---|---|
| Having your papers read and downloaded is... | Career: 3.90<br>Science: 3.90<br>Satisfaction: 4.10 | (SA) $F(2, 250) = 4.873$<br>$p = 0.008$ | 95% CI(−0.158, 0.158)<br>p = 1.000<br>md = 0.000 | **95% CI(−0.365, −0.048)**<br>**p = 0.011**<br>**md = −0.206** | **95% CI(−0.343, −0.070)**<br>**p = 0.003**<br>**md = −0.206** |
| Having public outreach (e.g., social media, news, etc.) is... | Career: 3.84<br>Science: 3.77<br>Satisfaction: 3.72 | (SA) $F(2, 250) = 1.251$<br>$p = 0.288$ | ——— | ——— | ——— |
| Having your results used or implemented in practice is... | Career: 4.02<br>Science: 4.26<br>Satisfaction: 4.37 | (SA) $F(2, 250) = 12.875$<br>$p < 0.001$ | **95% CI(−0.380, −0.096)**<br>**p = 0.001**<br>**md = −0.238** | **95% CI(−0.479, −0.203)**<br>**p < 0.001**<br>**md = −0.341** | 95% CI(−0.233, 0.027)<br>p = 0.118<br>md = −0.103 |
| Having luck is... | Career: 4.27<br>Science: 4.02<br>Satisfaction: 3.89 | (SA) $F(2, 250) = 14.229$<br>$p < 0.001$ | **95% CI(0.115, 0.377)**<br>**p < 0.001**<br>**md = 0.246** | **95% CI(0.240, 0.522)**<br>**p < 0.001**<br>**md = 0.381** | 95% CI(−0.022, 0.292)<br>p = 0.091<br>md = 0.135 |

* We report Greenhouse-Geisser (GG) tests when the results of Mauchly's test of sphericity could not confirm the sphericity of the data. Otherwise, Sphericity Assumed (SA) tests are reported. Abbreviations: confidence intervals (CI), mean difference (md)
